# Zika virus replication in the mosquito *Culex quinquefasciatus* in Brazil

**DOI:** 10.1101/073197

**Authors:** D. R. D. Guedes, M. H. S. Paiva, M. M. A. Donato, P. P. Barbosa, L. Krokovsky, S. W. dos S. Rocha, K. L. A. Saraiva, M. M. Crespo, R. M. R. Barbosa, C. M. F. Oliveira, M. A. V. Melo-Santos, L. Pena, M. T. Cordeiro, R. F. de O. França, A. L. S, de Oliveira, W. S. Leal, C. A. Peixoto, C. F. J. Ayres

**Affiliations:** Department of Entomology, Centro de Pesquisas Aggeu Magalhães, Fundação Oswaldo Cruz-Pernambuco. Av. Moraes Rego, s/n campus da UFPE, Cidade Universitária, Recife-PE, Brazil, CEP: 50670-420.; Universidade Federal de Pernambuco, Centro Acadêmico do Agreste - Rodovia BR-104, km 59 - Nova Caruaru, Caruaru - PE – Brazil, CEP: 55002-970.; Laboratory of Virology and Experimental Therapy (LAVITE), Centro de Pesquisas Aggeu Magalhães, Fundação Oswaldo Cruz-Pernambuco. Av. Moraes Rego, s/n campus da UFPE, Cidade Universitária, Recife-PE, Brazil, CEP: 50670-420.; Center for Statistics and Geoprocessing, Centro de Pesquisas Aggeu Magalhães, Fundação Oswaldo Cruz-Pernambuco. Av. Moraes Rego, s/n campus da UFPE, Cidade Universitária, Recife-PE, Brazil, CEP: 50670-420.; Department of Molecular and Cellular Biology, University of California-Davis, Davis, CA, 95616 USA.

**Author notes:** These authors contributed equally to this work (D. R. D. Guedes, M. H. S. Paiva).

**Keywords:** Zika, microcephaly, *Culex*, *Aedes*, vectorial competence, vector control

## Abstract

Zika virus (ZIKV) is a flavivirus that has recently been associated with increased incidence of neonatal microcephaly and other neurological disorders. The virus is primarily transmitted by mosquito bite, although other routes of infection have been implicated in some cases. The *Aedes aegypti* mosquito is considered to be the main vector to humans worldwide, but there is evidence of other mosquito species, including *Culex quinquefasciatus*, playing a role in the Brazilian outbreak. To test this hypothesis, we experimentally compared the vectorial competence of laboratory-reared *A. aegypti* and *C. quinquefasciatus*. We found ZIKV in the midgut, salivary glands, and saliva of artificially fed *C. quinquefasciatus*. Additionally, we collected ZIKV-infected *C. quinquefasciatus* from urban areas of high microcephaly incidence in Recife, Brazil. Take into account; these findings indicate that there may be a wider range of vectors for ZIKV than anticipated.

---

Zika is classically considered a mild disease whose symptoms include fever, joint pain, rash and, in some cases, conjunctivitis (*1*). However, the Zika outbreak in Brazil has been associated with an increased incidence of neonatal microcephaly and neurological disorders (*2*, *3*). Zika virus (ZIKV) is a poorly known, small, enveloped RNA virus with ssRNA (+) belonging to the Family *Flaviviridae.* It was first isolated in April 1947 from a rhesus monkey and in January 1948 from the mosquito species *Aedes africanus* (*4*). Since then, several ZIKV strains have been isolated from many samples, mostly mosquitoes, including species from the genera *Aedes*, *Mansonia, Anopheles* and *Culex* (*5*).

The first known Zika epidemic in an urban environment occurred in Micronesia in 2007, with approximately 73% of the human population on Yap island becoming infected (*6*). Intriguingly, although many *Aedes* mosquitoes were collected in the field and evaluated for virus detection, no samples were found to be positive for ZIKV (*6*). Additionally, it is important to highlight that *Aedes aegypti* (*A. aegypti*) is absent from most islands in the Micronesia archipelago and is very rare on the islands where it is present (*6*, *7*).

There is a global consensus among scientists and health agencies that *Aedes* spp. are the main ZIKV vector in urban areas (WHO, 2016). This is in part because vector competence experiments for ZIKV have been conducted exclusively for species of this genus, mainly *A. aegypti* (*8*, *9*). Previous laboratory studies (*8*, *10*) suggested that *A. aegypti* is a ZIKV vector. Recently, high rates of dissemination and transmission of the ZIKV in *A. aegypti* has been observed under laboratory conditions (*11*). Intriguingly, a few studies show that *A. aegypti* and Aedes albopictus populations have low rates of ZIKV transmission (*12*) or none (*13*, *14*), but the role of other vectors in the spread of ZIKV has been overlooked. Thus, other mosquito species, co-existing with A. aegypti in urban areas, could contribute to ZIKV transmission (*15*). Here, we report data that support the idea that *Culex quinquefasciatus*, the most common mosquito in tropical and subtropical areas, is a potential ZIKV vector. We performed mosquito vector competence assays under laboratory conditions, comparing both *A. aegypti* and *C. quinquefasciatus* using different virus doses, as well as the detection of ZIKV in wild *C. quinquefasciatus* mosquitoes. ZIKV was detected in salivary glands and in the saliva of artificially fed *C. quinquefasciatus* mosquitoes, suggesting that this species is a potential vector for ZIKV transmission. Taken together, our results have implications for vector control strategies and understanding the epidemiology of ZIKV.

## Vector competence assays

Artificial blood feeding assays were performed using two laboratory-reared colonies: RecLab (*A. aegypti*) and CqSLab (*C. quinquefasciatus*) and a field-collected population of *A. aegypti*(F_1_/F_2_) from the Archipelago of Fernando de Noronha, a district from Pernambuco state, Northeast Brazil. A local Zika virus strain, isolated from the serum of a patient with Zika symptoms from Pernambuco State, Brazil, during the 2015 outbreak (ZIKV/H.sapiens/Brazil/PE243/201), fully characterized (accession number KX197192.1), was used in vector competence assays.

Seven to ten days-old females were challenged in an artificial feeding, consisted of a Petri dish covered with Parafilm M^®^, with a mixture containing equal volumes of defibrinated rabbit blood and the viral suspension. Here, in each assay, we used two different viral doses: 10^6^ PFU/ml and 10^4^ PFU/ml. Mosquitoes were exposed for 90 minutes and, after that, only the engorged females were transferred to another cage and maintained in the infectory room, under bio safety conditions (BSL2) for 15 days. At 3, 7, 11 and 15 days post infection (dpi), midguts and salivary glands were dissected individually and transferred to 1.5 ml vials containing a mosquito diluent (20% of fetal bovine serum in PBS with 50 μg/ml penicillin/streptomycin, 50 μg/ml gentamicin and 2.5 μg/ml fungizone) and stored at −80°C until further usage. After RNA extraction, samples were assayed by quantitative RT-PCR (RT-qPCR) and both Infection Rate (IR), which is the proportion of infected midguts, and the proportion of infected salivary glands (SR), which is the number of positive salivary glands divided by the total number of salivary glands tested, were calculated for each species on each dpi. All procedures are described in details in the Supplemental Materials.

A total of 289 mosquitoes were examined for ZIKV infection by RT-qPCR. Among these mosquitoes, 130 were *A. aegypti* RecLab, 60 were *A. aegypti* FN and 99 were *C. quinquefasciatus.* During the extrinsic incubation period, a high mortality rate was observed in the infected group a few days after blood feeding, with peak mortality observed between 3-5 dpi. In *A. aegypti* populations, the mortality rate ranged from 48% to 52%; in *C. quinquefasciatus* mosquitoes, it ranged from 33% to 44% (data not shown).

In both species, we detected ZIKV in the midgut at most time-points under study, except in field-collected *A. aegypti* (FN) blood fed at a low viral dose (10^4^ PFU/ml). In the salivary glands of *A. aegypti*, we detected ZIKV-positive samples at 3 dpi for both viral doses. Although variations in infection were observed, the IR between *Aedes* and *Culex* mosquito’s species was not statistically significant (Fig 1A). Analysing only *Culex* species, the differences in IR were not statistically significant comparing different viral doses (p>0.05). When a high viral dose (10^6^ PFU/ml) was used during artificial feeding, the SR reached an average of 60% in *A. aegypti* RecLab, and 100% in *C. quinquefasciatus* at 7 dpi, but this average declined to 20% at 15 dpi in *Culex* and to 50% in *A. aegypti*. However, this difference was not significant (p>0.05) (Fig 1B). When a lower viral dose was used, for both *A. aegypti* populations (lab and field-caught), we observed that the mosquito lab colony infection rate was higher than field-caught mosquitoes (Figure 1A p=0.0186), although there were no differences in SR (Figure 1B p>0.05). The maximum SR in *Culex* at lower viral dose, was 10% at 3dpi declining to zero after 15 dpi (p>0.05) (Fig. 1B). We also sampled mosquitoes at 11 dpi in the first trial, except for field-collected *A. aegypti* (FN), which had a SR of 10%; neither *C. quinquefasciatus* nor *A. aegypti* samples from the lab were ZIKV positive (data no shown).

**Fig. 1:**
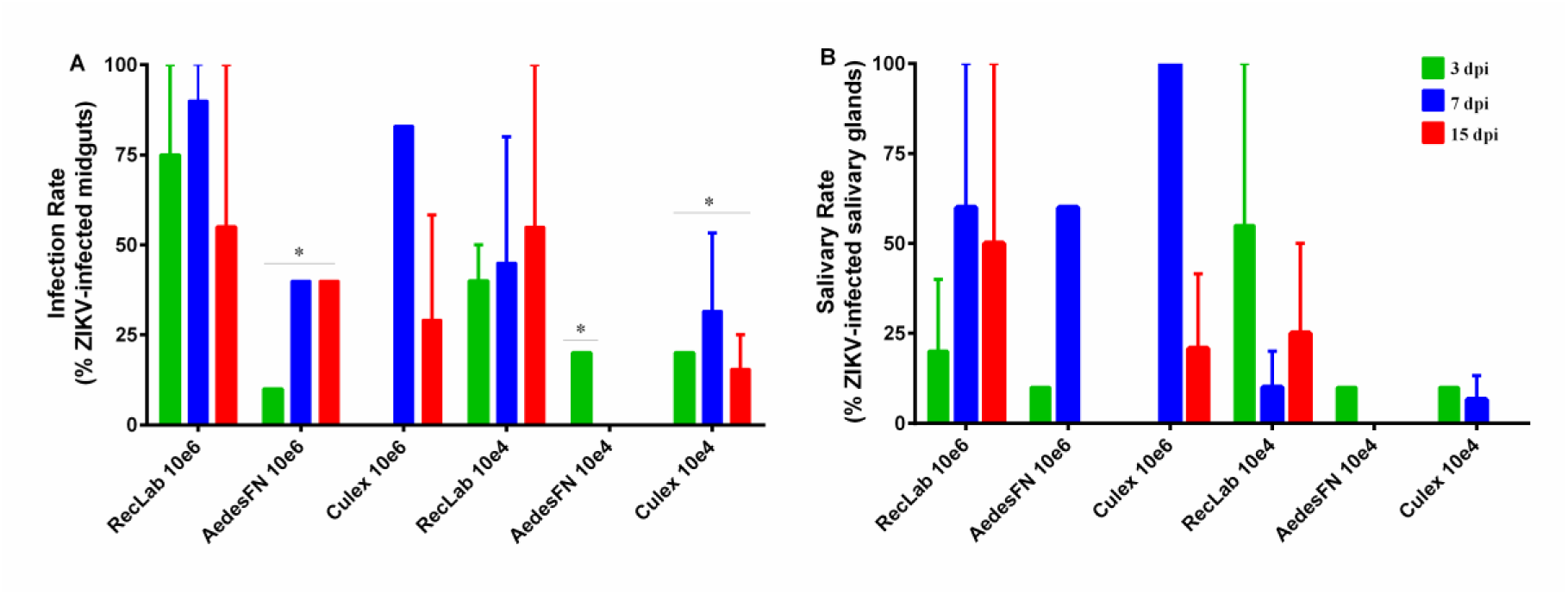
Experimental infection of ZIKV in laboratory-reared *A. aegypti* and *C. quinquefasciatus* collected 3 (green bars), 7 (blue bars) and 15 days post infection (dpi) (red bars). (A) Proportions of ZIKV-positive midguts at each sampling point (average mosquitoes per group = 10 for two replicates). (B) Proportions of ZIKV-positive salivary glands at each sampling point (average of mosquitoes per group = 10 for two replicates). Significance was determined using one-way ANOVA with Tukey’s multiple comparison test (* p < 0.05).

RT-qPCR was used to quantify ZIKV RNA load at the different time-points. In general, viral RNA copies in *A. aegypti* RecLab in the midguts and salivary glands varied considerably. Both *A. aegypti* FN and *C. quinquefasciatus* viral copies in target organs (midgut and salivary glands) remained detectable (Fig. 2A to D). To evaluate ZIKV transmission in saliva for both species, honey-soaked filter papers (FTA Classic Cards, Whatman^®^, Maidstone, UK) were offered to mosquitoes to feed upon 8-14 dpi. At 9-12 dpi, ZIKV RNA was detected in saliva of both *A. aegypti* and *C. quinquefasciatus* species (Fig. 3). When a high viral dose (10^6^) was used, the amount of viral RNA copies expectorated during salivation in both Aedes and Culex were similar at all time-points analysed (p>0.05). However, when the mosquitoes were challenged with a low viral dose (10^4^), *A. aegypti* expectorated more RNA viral copies than *Culex* (p=0.0473).

**Fig. 2:**
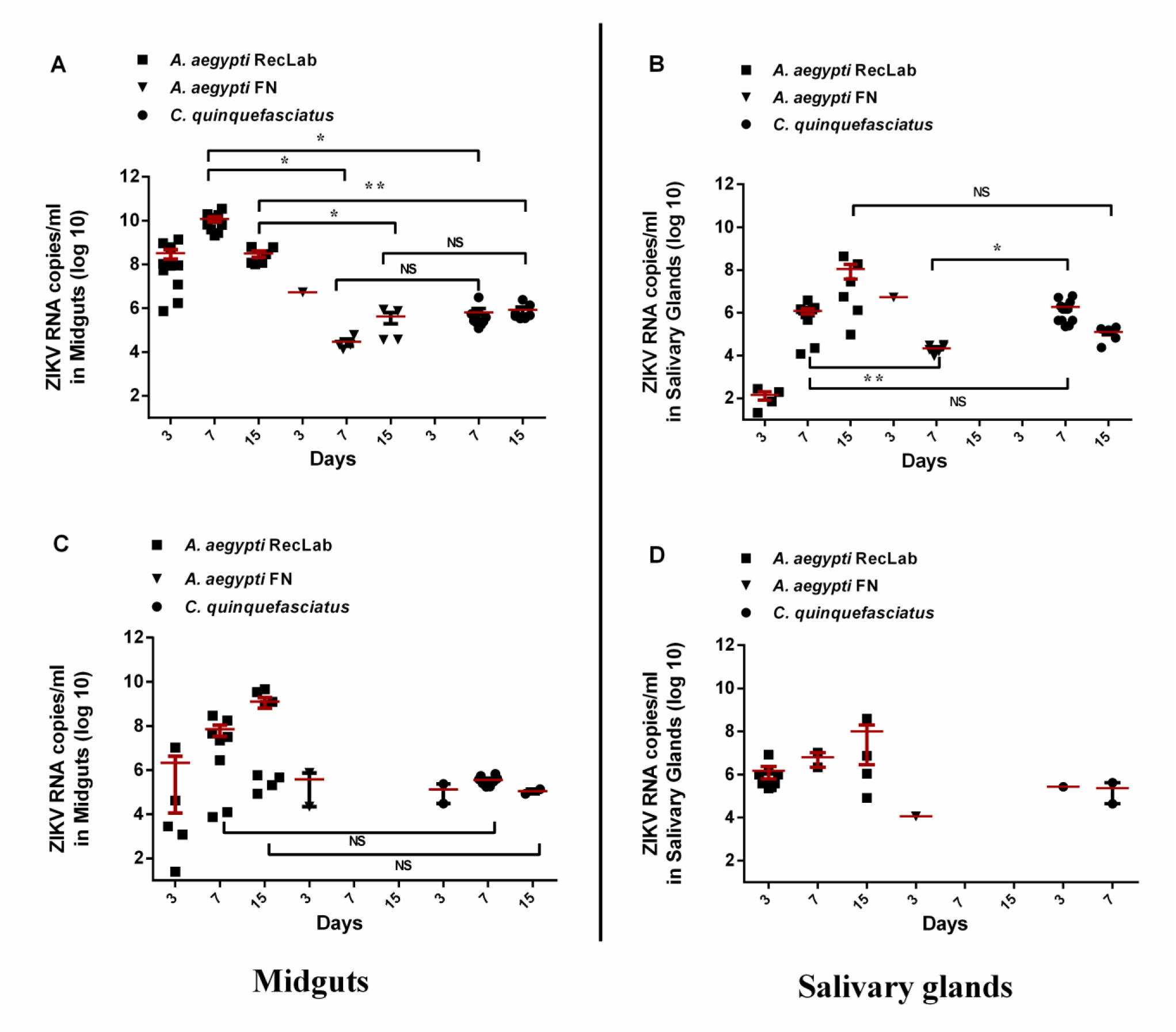
Quantification of RNA viral copy number in the midguts and salivary glands of *A. aegypti* and *C. quinquefasciatus* mosquitoes experimentally fed with blood containing ZIKV at 10^6^ PFU/ml (A, B) and 10^4^ PFU/ml (C, D). Squares represent *A. aegypti*(RecLab) population, inverted triangles represent *A. aegypti*(FN) population and circles represent *C. quinquefasciatus*. Significance is shown in the bars and was determined using an unpaired t-test (* p < 0.05, ** p < 0.01).

**Fig. 3:**
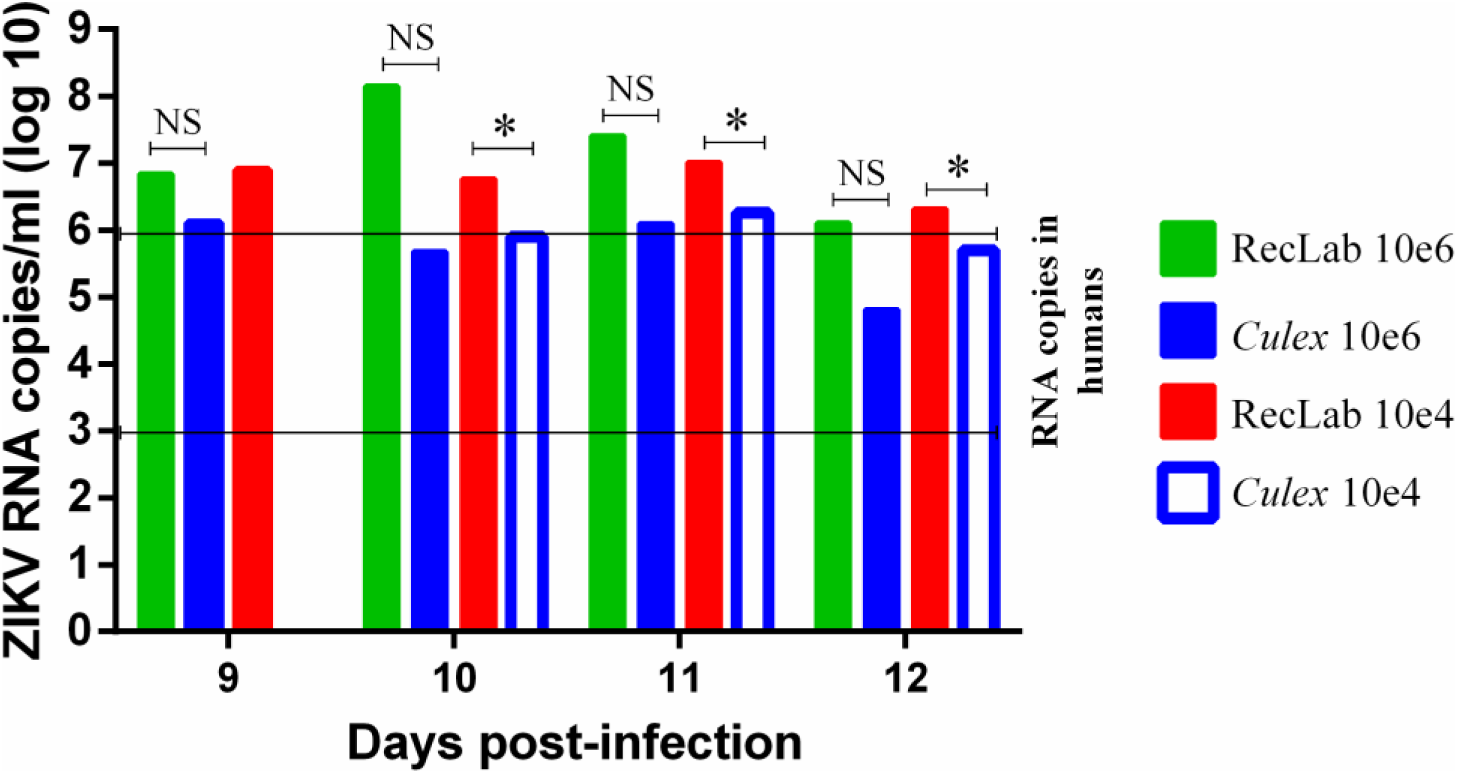
Quantification of ZIKV in *A. aegypti* and *C. quinquefasciatus* saliva expectorated onto FTA cards 9 - 12 days post infection (dpi). Green bars show *A. aegypti*(RecLab) population blood-fed with ZIKV at 10^6^ PFU/ml, solid blue bars show *C. quinquefasciatus* population blood-fed with ZIKV at 10^6^ PFU/ml, red bars show *A. aegypti*(RecLab) population blood-fed with ZIKV at 10^4^ PFU/ml and open blue bars show *C. quinquefasciatus* population blood-fed with ZIKV at 10^4^ PFU/ml. Significance was determined by an unpaired t-test (* p < 0.05).

## Transmission Electron Microscopy

To further confirm our results from RT-qPCR, we performed a transmission electron microscopy from dissected salivary glands from *C. quinquesfaciatus* infected mosquitoes. The morphological organization of *C. quinquefasciatus* salivary glands showed an electron-dense apical cavity, displaying membrane projections extending from the wall (Fig. 4A, B). ZIKV infected salivary acinar cells of *C. quinquefasciatus* showed signs of cytopathic disruptions, including cisternae in the endoplasmic reticulum and tubular proliferated membranes, organized in several patches within a single cell (Fig. 4C, D). Mature ZIKV particles of 40-50 nm in diameter, composed of a central electrodense core (∼30 nm in diameter) surrounded by a viral envelope, were observed inside the dilated endoplasmic reticulum (Fig. 5A to D). In some regions, viral envelope formation is shown to arise from endoplasmic membrane (Fig. 5B). Some ZIKV virions were observed proximal to the apical cavity of the salivary cell. Mitochondria also showed severe damage, including complete loss of cristae (Fig. 5D). In summary, transmission electron microscopy analysis confirmed that *C. quinquefasciatus* mosquitoes are permissive to ZIKV infection, since viral particles were detected at the salivary glands of artificially fed mosquitoes.

**Fig. 4:**
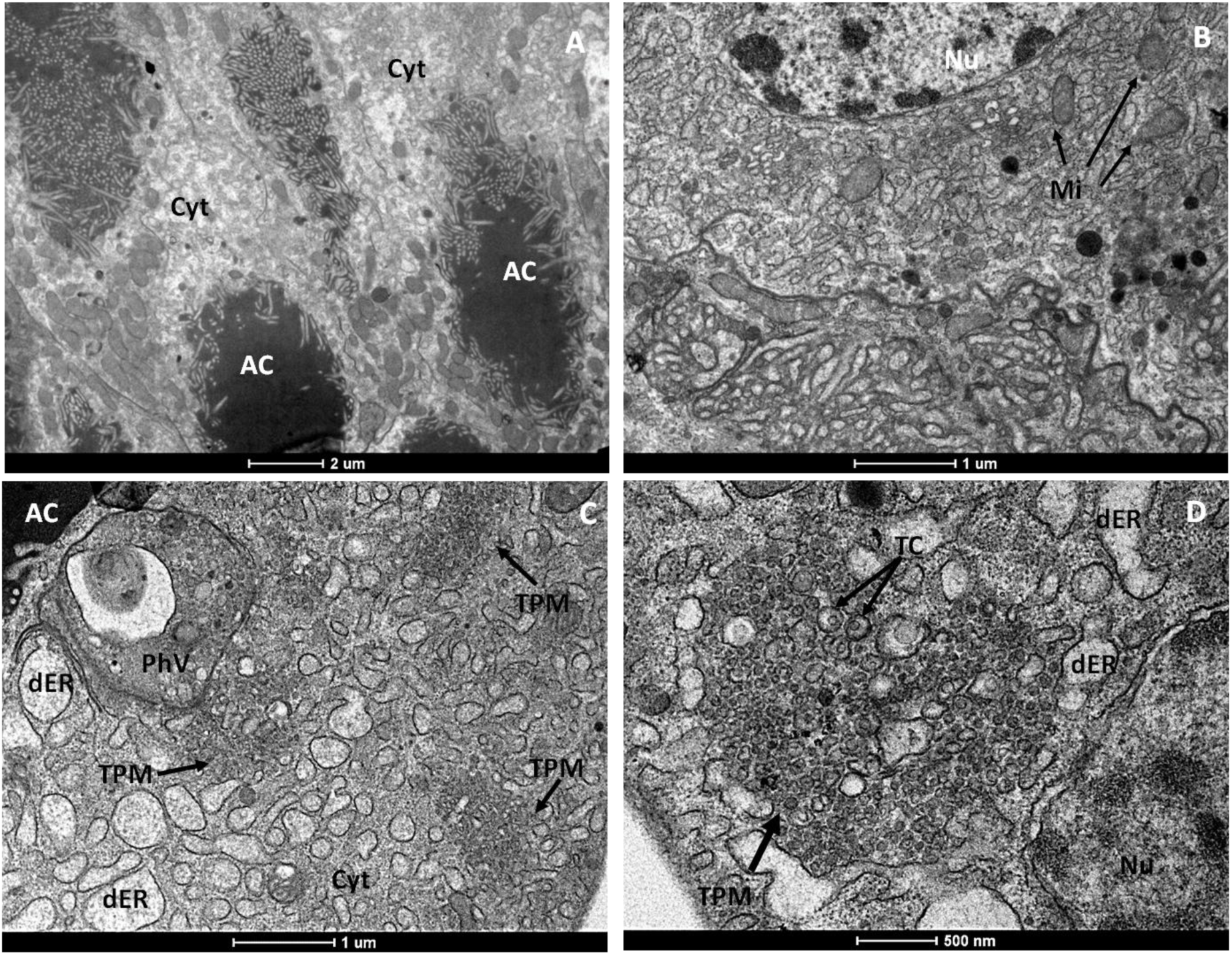
(**A-B**) Ultrathin sections of uninfected *C. quinquefasciatus* salivary gland. (**A**) Shows the electrodense content of the apical cavity (AC) with membrane projections extended from the wall. (**B**) Uninfected acinar salivary gland cell showing Nu, nucleus; Cyt, cytoplasm; ER, endoplasmic reticulum; Mi, mitochondria. (**C-D**) Cytopathic effects of salivary glands cells infected with ZIKV showing several patches of tubular proliferated membrane (TPM), distended endoplasmic reticulum (dER) and a phagolysozome-like vacuole (PhV). Cyt, cell cytoplasm; TC; thread-like centers.

**Fig. 5:**
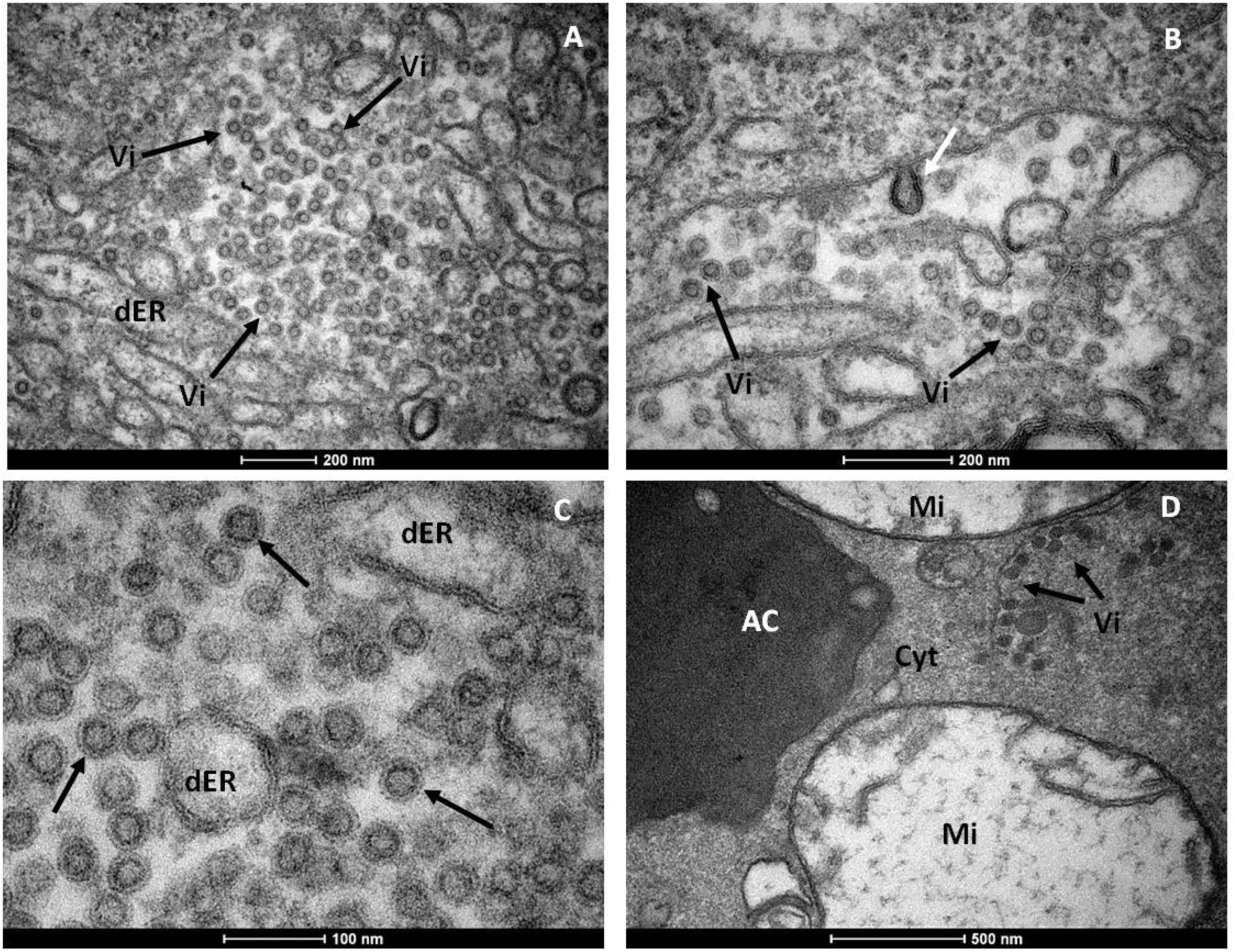
Mature ZIKV particles inside *C. quinquefasciatus* salivary gland cell. (**A**) Numerous ZIKV within dilated endoplasmic reticulum (dER). (**B**) Envelope formation from endoplasmic membrane (white arrow). (**C**) Showing enveloped virus particles with electrodense cores. (**D**) Viral particles accumulated proximal the acinar cavity (arrows), note damaged mitochondria. Cyt, cell cytoplasm; AC, acinar cavity, Mi, mitochondrion; Vi, virion (s).

## ZIKV detection in field-caught *C. quinquefasciatus*

Lastly, we conducted ZIKV surveillance (February to May 2016) with mosquitoes collected with a battery-operated aspirator device (Horst^®^) from residences inhabited by individuals with clinical symptoms of zika fever. Field-collected mosquitoes were sorted by place of collection, species, sex, feeding status (engorged and not engorged) and grouped in pools of up to 10 mosquitoes. A total of 1,496 adult *C. quinquefasciatus* and 408 *A. aegypti* female mosquitoes were collected from different sites in the Metropolitan Region of Recife (Fig. S1). These mosquito pools were grinded in Leibovitz medium supplemented with 5% FBS. These samples were separated into two aliquots, one for RT-qPCR and the other for virus isolation. From 270 pooled-samples of adult female *C. quinquefasciatus* and 117 pools of *A. aegypti* mosquitoes assayed by RT-qPCR, three *Culex* and two *Aedes* pools were positive for ZIKV. Interestingly, two out of the three positive *Culex* samples were not blood-fed, whereas concerning *Aedes* pools, the two positive pools for ZIKV were fed. The cycle threshold (Ct) of *Culex* positive pools when screened by RT-qPCR were 37.6 (sample 5), 38.0 (sample 17) and 38.15 (sample 163). Concerning *Aedes* pools, Cts were 37.5 (sample 3) and 37.9 (sample 7). Minimum infection rate (MIR - number of positive pools divided by the total of specimens assayed multiplied by 1000) were calculated for both species. For *C. quinquefasciatus*, MIR was 2.0 and concerning *A. aegypti*, MIR was 4.9. In an attempt to isolate ZIKV from field-caught *Culex* mosquitoes, we inoculated in African Green Monkey kidney cells, samples from two positive pools with the lowest cycle thresholds. Indeed, ZIKV was isolated from these samples, thus unambiguously demonstrating that this species was carrying active ZIKV particles in Recife, Brazil. Two ZIKV-positive isolates from field-caught *Culex* mosquitoes were submitted to Sanger and MinION platforms. Sanger sequencing resulted in low quality sequences and only a partial fragment was acquired from MinION sequencing, probably as a direct result of low viral titers. This partial sequence enabled us to confirm the virus identity. Sequence was deposited at GenBank, and accession number is still to be provided.

## Discussion

Our work has associated a second mosquito genus in ZIKV transmission cycle in Northeastern Brazil. We showed that, *C. quinquefasciatus*, also known as the southern house mosquito, which is the most common mosquito in urban areas in Brazil, is susceptible to infection with ZIKV during experimental blood feeding; moreover, we found that ZIKV has an active replication cycle in the salivary glands and being subsequently released in the saliva. In addition, we were able to detect ZIKV circulating in wild *C. quinquefasciatus* collected from Recife.

Although it is widely assumed that *A. aegypti* is the main ZIKV vector, previous vector competence studies are inconclusive. In the present study, a low dose of ZIKV (10^4^ PFU/ml) was used for comparison with the higher doses used in previous studies (*11*, *12*, *14*). We found that both *A. aegypti* and *C. quinquefasciatus* can be experimentally infected by ZIKV even at low doses and that ZIKV virus was subsequently detected in the saliva.

Cornet et al. (*10*) concluded that not all infected mosquitoes could transmit the virus and could not always transmit it, in contrast to the idea that once infected, a mosquito would transmit virus for its entire life. This finding suggests that a time window for vector-borne ZIKV transmission may exist. We found that after 11 dpi, most samples were negative for ZIKV (apart from one positive salivary gland of *A. aegypti* given 10^6^ PFU/ml), thus our maximum time point analysis was set to 15 days post infection. However, Boorman & Porterfield (*8*) reported that virus replication resumed at 15-20 dpi and ZIKV remained present in *Aedes* mosquitoes for up to 60 days.

To confirm that the virus detected in the salivary glands by RT-qPCR was being released in saliva during consecutive blood meals, we followed up the viral load from days 8 to 14 post-infection using filter paper cards. This strategy of viral RNA detection directly from FTA cards has been employed in previous studies for arbovirus surveillance (*16*, *17*). In the present study, we successfully detected ZIKV RNA copies in cards from *A. aegypti* and *C. quinquefasciatus* populations. This result demonstrates that in addition to being susceptible to ZIKV infection, allowing virus replication in the salivary glands, both species are capable of effectively transmit ZIKV.

RT-qPCR results were confirmed by transmission electron microscopy. The general mature ZIKV morphology observed on the salivary glands confirmed previous ultrastructural studies (*18*-*20*). In salivary glands cells, ZIKV replication causes cytopathic effects by 7 dpi. Similar results have been shown for West Nile virus (WNV) (*21*, *22*), although we did not directly observe ZIKV nucleocapsids budding from endoplasmic reticulum membranes nor from the tubular proliferated membrane (*21*). The fact that we found salivary glands positive for ZIKV when the midgut of the same mosquito was negative indicates that mosquitoes may be clearing viral infection in the midgut while virus replication continues in the salivary glands. This finding has implications for the analytical methods employed in vector competence studies.

Currently, there is a lack of studies investigating *Culex* vectorial competence for ZIKV. Most studies have targeted only *Aedes* species, and only a few studies have compared different species (including *Culex*) regarding natural infection rates. Surprisingly, Diallo et al. (*5*) observed a higher minimum infection rate for *Culex perfuscus* (10x higher) than for *A. aegypti*. Positive *A. aegypti* samples have always been reported at very low infection rates, even in areas with high human ZIKV infection rates, such as Malaysia (*23*). Indeed, in Micronesia (*6*), and French Polynesia, ZIKV was not detected in wild-caught *Aedes* spp. mosquitoes during outbreaks. It is interesting that in all of these areas, *C. quinquefasciatus* is an abundant mosquito species that may have also played an undetected role in ZIKV transmission. Furthermore, *A. aegypti* and *C. quinquefasciatus* have completely different behaviours regarding feeding periods and breeding site preferences.

Thus, our findings indicate that vector control strategies may need to be re-examined since reducing *A. aegypti* populations may not lead to an overall reduction in ZIKV transmission if *Culex* populations are slight affected by *Aedes* specific control measures. To the moment, there is no broad ongoing program for *C. quinquefasciatus* control in Brazil, although Recife, Olinda and Jaboatão dos Guararapes, three municipalities in Recife Metropolitan Region, have undertaken specific control of *C. quinquefasciatus* to control lymphatic filariasis transmission locally (*24*).

Viral transmission via *C. quinquefasciatus* is not a new concept; this species is the major vector of West Nile virus in North America (*25*), along with Japanese encephalitis virus (*26*) and equine encephalitis virus (*27*). Our present study indicates that *C. quinquefasciatus* mosquitoes may be involved in ZIKV transmission in Recife. Thus, it is now necessary to understand the contributions of each species in transmission to target each one properly. In conclusion, considering its high abundance in urban environments and its anthropophilic behaviour in Brazil (*28*-*30*), *C. quinquefasciatus* may be a vector for ZIKV in this region.

## Acknowledgements

This work was supported in part by the Fundação de Amparo à Pesquisa do Estado de Pernambuco (FACEPE; APQ-1608-2.13/15 and APQ-0085-2.13/16 to C.F.J.A.) and the National Institute of Allergy and Infectious Diseases of the National Institutes of Health (R01AI095514 and 1R21AI128931-01 to W.S.L.) C.F.J.A. and C.A.P. are supported by productivity fellowship from the Brazilian National Council for Research and Development (CNPq). We thank the staff of the insectary at Aggeu Magalhães Research Center for technical assistance, the Program for Technological Development in Tools for Health (PDTIS-FIOCRUZ) for allowing us to use their facilities, and the staff of the Pernambuco State Health Department for sharing recent data on microcephaly and assisting in surveillance.

## Supplementary Materials for

### Material and Methods

#### Mosquitoes

The present study was conducted using two laboratory colonies of mosquitoes and field-collected specimens of *Aedes aegypti* (F_1_-F_2_) from the Archipelago of Fernando de Noronha, a district of Pernambuco state (PE). *Culex quinquefasciatus* (formerly known as *C. pipiens quinquefasciatus*) mosquitoes originated from eggs (rafts) collected in Peixinhos, a neighborhood from Recife, PE, Brazil in 2009. This colony (CqSLab) was founded with approximatelly 500 field collected rafts (about 200,000 mosquitoes), and analysis of 16 microsatelitte loci showed that levels of genetic variation found in the CqSLab was similar to those found in field-caught mosquitoes (*31*). The *A. aegypti* laboratory colony (RecLab) was established with approximately 1,000 specimens collected in Graças, a neighborhood from Recife Metropolitan Region and maintained in the insectary of CPqAM/FIOCRUZ since 1996, under standard conditions: 26 ± 2°C, 65–85% relative humidity, 12/12 light/dark cycle. More information regarding the two laboratory colonies are described elsewhere (*32*, *33*). The mosquitoes were kept in the insectary of the Department of Entomology, FIOCRUZ/PE, under standard conditions described above. Larvae were maintained in plastic trays filled with potable water and were fed solely on cat food (Friskies^®^), while adults were given access to a 10% sucrose solution until they were administered defibrinated rabbit blood infected with the ZIKV.

#### Virus strain

Experimental infections of mosquitoes with ZIKV were conducted using the ZIKV BRPE243/2015 strain derived from the serum of a patient with an acute maculopapular rash in Pernambuco State, Brazil, during the 2015 outbreak. This strain was named ZIKV/*H. sapiens*/Brazil/PE243/2015 strain, according to the nomenclature described by Scheuermann (*34*). Following isolation, the virus was passed once on *A. albopictus* C6/36 cells. Viral stocks were then produced in Vero cells and stored at −80°C until use. Prior to oral infection, the stock viral titer was calculated via plaque assay on Vero cells and reached 10^6^ plaque-forming units per millilitre (PFU/mL).

#### Artificial feeding

We conducted two artificial-feeding assays using a viral stock concentration of 10^6^ PFU/ml and a 100-fold diluted viral stock. Of note, in the first artificial feeding assay, frozen virus sample was mixed with defibrinated rabbit blood. In the second assay, ZIKV BRPE243 was first grown in Vero cells at a multiplicity of infection (MOI) of 0.5 for 4-5 days. Subsequently, the cell culture flasks were frozen at −80°C, thawed at 37°C twice, and then mixed with defibrinated rabbit blood in a 1:1 proportion. Seven- to tenday-old female mosquitoes that were under a 10% sugar solution were starved for 18 hours prior to artificial feeding. Mosquitoes were exposed to an infectious blood meal for 90 minutes, as described in Salazar *et al.* (*35*), with minor modifications. Briefly, infectious blood was provided in a membrane-feeding device, placed on each mosquito cage. The blood feeding was maintained at 37°C, by using heat packs during the process. Fully engorged mosquito females were cold anesthetized, transferred to a new cage and maintained in the infection room for 15 days. Both assays included a control group fed on uninfected culture cells mixed with defibrinated rabbit blood. After the infectious blood meal was administered, the mortality rate was estimated daily for each cage, including that of the control group. Dead mosquitoes were removed from the cages.

#### RNA extraction and virus detection

Four to fifteen mosquitoes were dissected in order to collect midgut and salivary glands at 3, 7 and 15 dpi. Tissues were individually transferred to a 1.5 ml DNAse/RNAse free microtube containing 300/l of mosquito diluent (*36*) and stored at −80°C until further usage. Each tissue was ground with sterile micropestles and RNA extraction was performed with 100 µl of the homogenate. The TRizol^®^ method (Invitrogen, Waltham, MA) was performed, according to the manufacturer’s instructions with modifications as follows. Tissues homogenate (100 sl) was mixed with 200 el of Trizol, homogenized by vortexing for 15 seconds and incubated for 5 minutes at room temperature. Chloroform (100 ll) was added to the mixture and the homogenization was performed by shaking tubes vigorously for 15 seconds by hand. Mixture was then incubated at room temperature for 2-3 minutes. Samples were centrifuged at 12,000 x g for 15 minutes at 4°C. Aqueous phase of each sample was removed and transferred to a new tube containing 250 Cl of 100% isopropanol. Mixture was incubated at room temperature for 10 minutes and then centrifuged at 12,000 x g for 10 minutes at 4°C. Supernatant was removed and RNA pellet was washed with 300 Cl of 75% ethanol. Samples were homogenized briefly then centrifuged at 7,500 x g for 5 minutes at 4°C. Supernatant was discarded and RNA was then air dried for 15 minutes. RNA pellet was resuspended in 30 nl of RNAse free water. After RNA resuspension, samples were treated with DNAse (Turbo^TM^ DNase, Ambion) according to the manufacture’s protocol.

Virus detection was performed by quantitative RT-PCR (RT-qPCR) in an ABI Prism 7500 SDS Real-Time system (Applied BioSystems, Foster City, CA, USA), using the QuantiNova Probe RT-PCR kit (Qiagen, Hilden, Germany). RT-qPCR was performed in a 20 µl reaction volume containing 5 µl of extracted RNA, 2x QuantiNova Probe RTPCR Master Mix, 0.2 µl QuantiNova Probe RT Mix, 0.1 µl ROX Reference Dye, 100 µM of each primer (stock) and 25 µM of the probe (stock). Primers, probe and PCR conditions were first described in Lanciotti *et al.* (*37*) and each sample tested in duplicates. RT-qPCR cycling followed a single cycle of reverse transcription for 15 minutes at 45°C, 5 minutes at 95°C for reverse transcriptase inactivation and DNA polymerase activation, and then 45 cycles of 5 seconds at 95°C and of 45 seconds at 60°C (annealing-extension step). The amount of viral RNA from each sample was estimated by the comparison of cycle threshold values (Ct) to the standard curve for every RT-qPCR assay. The standard curve consisted of different dilutions of previously titrated ZIKV BRPE243/2015 RNA. Mosquitoes collected immediately after artificial feeding were used as positive controls, while control mosquitoes fed on uninfective blood and RT-PCR reactions containing no RNA represented negative controls. Fluorescence was analyzed at the end of the amplifications. Positive samples were used to calculate vector competence parameters, such as: infection rate (IR) which is the number of positive midguts divided by the total number of midgut tested; and proportion of infected salivary glands (SR), which is the number of positive salivary glands divided by the total number of salivary glands tested.

#### Collection of virus-infected mosquito saliva

To confirm if the virus detected by RT-qPCR within the salivary glands could be released during blood feeding meals, we assayed ZIKV in saliva samples. During 8-14 dpi, mosquitoes from each group were exposed to honey-soaked FTA Classic Cards (Whatman^®^, Maidstone, UK) to collect mosquito saliva. Each FTA card was prepared in a sterilized Petri dish and soaked in approximately 10 g of anti-bacterial honey (Manuka Honey Blend, Arataki Honey Ltd, New Zealand) and 1 ml of blue food dye for 2 hours. The blue food dye was used to determine if the mosquitoes had fed on the FTA cards. After 24 hours of exposure, each card was placed in a 15 ml falcon tube and stored at – 80°C until further use. To extract the RNA, cards were individually placed in a 2 ml microtube with 600 Cl of UltraPure^TM^ DNase/RNase-Free Distilled Water (ThermoFisher Scientific^®^, Massachusetts, USA). These eluted samples were kept on ice and vortexed 5 times for 10 seconds each. This process was repeated for 20 minutes. RNA was recovered from the FTA cards using the TRIzol^®^ method and was used to detect ZIKV, as described previously.

#### Transmission Electron Microscopy

Salivary glands of *C. quinquefasciatus* were dissected on 7 dpi, fixed for 2 hours in a solution containing 2.5% glutaraldehyde and 4% paraformaldehyde in 0.1 M cacodylate buffer solution. After fixation, the samples were washed twice in the same buffer and post-fixed in a solution containing 1% osmium tetroxide, 2 mM calcium chloride and 0.8% potassium ferricyanide in 0.1 M cacodylate buffer, pH 7.2, dehydrated in acetone as previously reported (*38*) and embedded in Fluka Epoxy Embedding kit (Fluka Chemie AG, Switzerland). Polymerization was performed at 60°C for 24 h. Ultrathin sections (70 nm) were placed on 300-mesh nickel grids, counterstained with 5% uranyl acetate and lead citrate, and examined using a transmission electron microscope (Tecnai Spirit Biotwin, FEI).

#### Statistical analysis

Infection rate (IR) and proportion of infected salivary glands (SR) were calculated for each species at different time points. Infection rate corresponded to the number of positive midguts divided by the total number of mosquitoes assayed. Proportion of infected salivary gland corresponded to the number of positive salivary glands divided by the total number of salivary glands assayed. Differences in both infection and transmission rates between species and viral load were analysed using GraphPad Prism^®^ software v.5.02. This software was used to plot graphics and to compare viral genome quantification values between different time-points, tissues and samples using an unpaired t-test and one-way ANOVA with Tukey’s multiple comparison tests.

#### ZIKV detection, viral isolation and sequencing in field collected mosquitoes

Mosquito samples were collected in the metropolitan region of Recife, from February to May 2016, in two distinct types of location: premises where zika cases were notified and at public Emergency Care Units (ECU). The Pernambuco Secretary of Health personnel carried out collections at ECUs, and our own fieldwork team collected mosquitoes in the premises. Both sets of collections were performed using a battery-operated aspirator (Horst^®^). Samples were sent alive to the Aggeu Magalhães Research Center (CPqAM), anesthetized at 4°C, morphologically identified, sorted by species, locality, sex, feeding status (engorged and not engorged), pooled (1–10 individuals/pool) and preserved at –80°C until assayed for RNA extraction and qRTPCR as described above. The minimum infection rate (MIR) for ZIKV in adults captured in the field was calculated as: (number of positive pools for ZIKV/total number of mosquitoes tested) x 1000 (*39*).

Positive samples were assayed for virus isolation in Vero cells as follows. In a tissue culture tube (TPP^®^), 1 ml of a 5 x 10^5^ cells/ml suspension in MEM medium were seeded for 24 h to form a monolayer. After that, the MEM medium was discarded and 1 ml of the filter homogenate (100 tl of positive homogenate + 900 ll of MEM medium) was inoculated in the cells. After 1h for virus adsorption, 1ml of fresh medium was added to the tissue culture tubes and they were then incubated at 37°C in 5% CO_2_ atmosphere until the detection of cytopathic effect. ZIKV positive samples collected from field mosquitoes were sequenced. Amplicons were generated by RT-PCR using Cloned AMV Reverse Transcriptase (Invitrogen, Waltham, MA). cDNAs were submitted to PCR reactions using primers targeting a 500 bp region spanning the capsid-envelope region of the virus (FW: 5’-CAATATGCTAAAACGCGGAGT-3 and REV: 5’-GGTTCCGTACACAACCCAAG-3’), under the following conditions: 94°C for 5 minutes, followed by 30 cycles of 94°C for 1 minute, 60°C for 1 minute and 72°C for 2 minutes, with a final extension of 72°C for 10 minutes. RT-PCR products were submitted to Sanger sequencing in an ABI 3500xL (Applied Biosystems, Carlsbad, CA). Sequences were edited and analyzed using CodonCode Aligner, v.3.7.1 (CodonCode Corporation, Dedham, MA). Both RNAs from ZIKV-positive isolates were also submitted to the NGS (Next-Generation Sequencing) platform: MinION (Nanopore, Oxford, UK). MinION sequencing was performed according to manufacturer’s protocol.

#### Spatial analysis

We georeferenced the points from where mosquitoes were collected using WGS-84 (World Geodesic System), a GPS receiver navigation and processed the data in a QGIS software. We generated a geographical database and performed a kernel density analysis based on the spatial distribution of reported cases of microcephaly registered by the Health Department of Pernambuco. The illustrated map shows an overlay between the location of the mosquito sampling, and the Kernel density map of reported cases of microcephaly from August 2015 to March 2016.

**Fig. S1:**
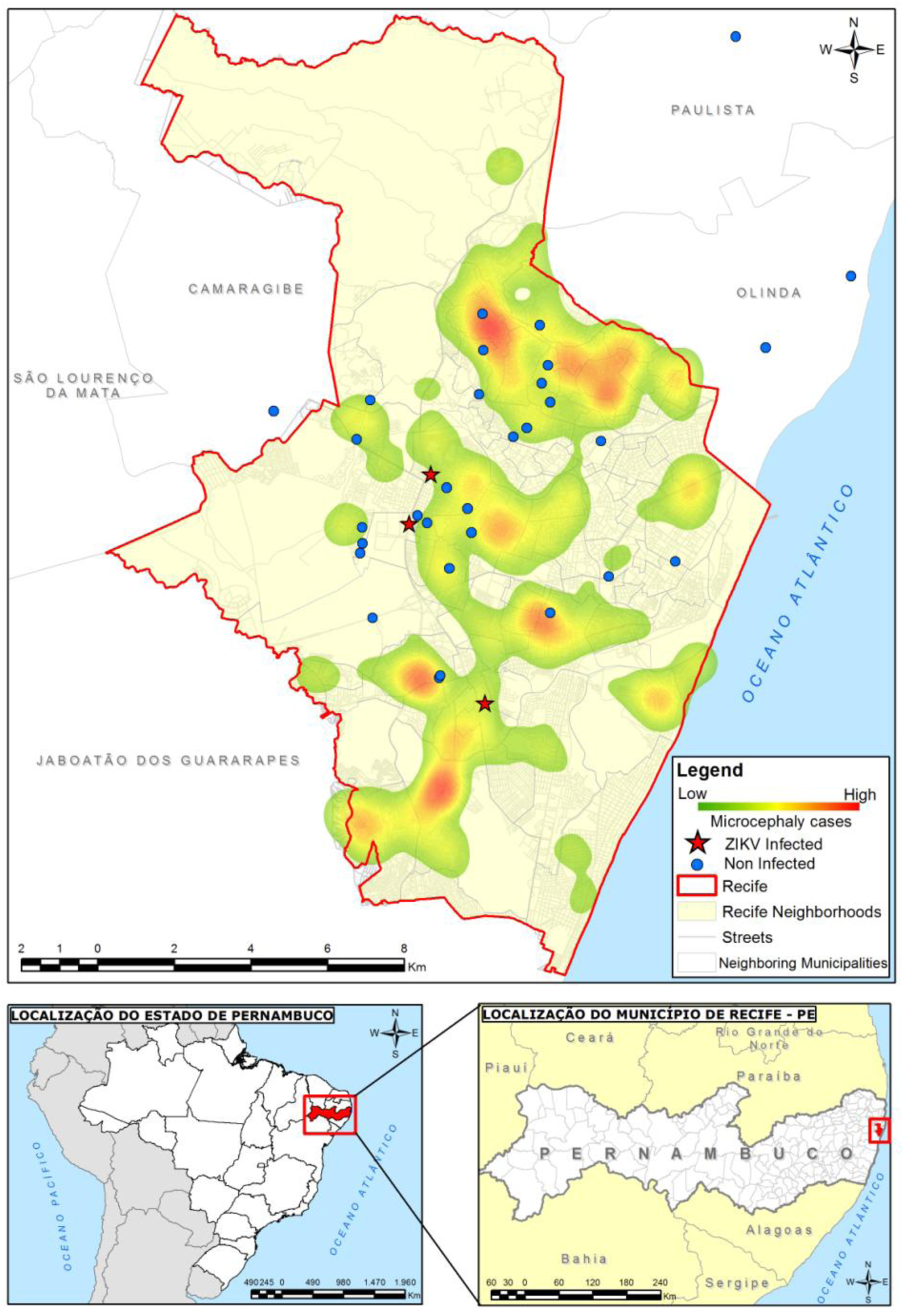
Kernel density map of microcephaly reported cases versus Point Map of the mosquito collection sites (with positive and negative *Culex* samples for the presence of ZIKV).

